# Evolutionary rescue in a consumer-resource system depends on the affected ecological traits and the population’s resident life-history

**DOI:** 10.1101/2025.08.04.668505

**Authors:** Asad Hasan, Michael C. Whitlock

## Abstract

With evolutionary rescue, a population that is declining due to an environmental change adapts to its environment, avoiding extinction. Previous theoretical work has explored the effects of negative density-dependence on rescue, showing that it may aid or hinder persistence. However, these models typically assume that it is only population intrinsic growth rates, *r*, or carrying capacity, *K*, that are negatively affected, and do not model density-dependence explicitly. Here, we analyze evolutionary rescue in a consumer-resource species following an abrupt environmental change, characterizing how rescue is dependent on the ecological effects of the environmental change on the consumer, which differently affect *r* and *K* through subsequent interactions with an explicit non-substitutable resource species. We derive approximate analytical predictions for the fixation probabilities of beneficial alleles, mutational supply, and times to extinction, which work well when selection is weak and individual turnover rates are low. We demonstrate that consumer rescue is dependent on the ecological effect of the environmental change, the resident life-history of the population, and the genetic architectures of evolving traits (monogenic versus polygenic). Our model suggests that measurements of intrinsic growth rates alone will be insufficient to predict rescue probabilities. This work extends our understanding of the interplay between ecology and evolution in influencing population persistence.

## Introduction

Understanding the factors underlying the ability of populations to persist in response to environmental change is of utmost importance, particularly in light of the rapidly accelerating loss of biodiversity due to climate change (Berg et al. 2010, Bellard et al. 2012). One such mechanism of persistence is through evolutionary adaptation, or specifically evolutionary rescue (see Carlson et al. 2014; Bell 2017 for reviews), the process by which a population that is declining due to an environmental perturbation subsequently evolves and adapts to its new environment, avoiding local extinction. Theoretical models of evolutionary rescue (hereafter ‘rescue’) are characterized by the concurrent modelling of both evolutionary and demographic processes, whereby successful rescue of a population is contingent on its rate of adaptation outpacing its rate of demographic decline (Gomulkiewicz & Holt 1995). Classical models of rescue have typically considered a single isolated species evolving to avoid extinction (e.g. Gomulkiewicz & Holt 1995; Burger & Lynch 1995; Lande & Shannon 1996). This work has recovered many important insights into the factors underlying persistence, including the positive roles of additive genetic variance in fitness and of initial population size. Later work has extended this theory to consider in detail the effects of population structure (Uecker et al. 2014), recombination (Uecker & Hermisson 2016), generation time (Draghi et al. 2024), and various other factors (Orr & Unckless 2008, 2014; Uecker 2017; Osmond et al. 2020; Xu 2023). The simplifying assumption of a single species existing in isolation, however, fails to capture ecological interactions among individuals such as predation and competition.

Ecological interactions may drastically influence demographic and evolutionary outcomes, and therefore rescue probabilities. Predation typically imposes a demographic cost on prey species, accelerating demographic decline. However, predation may also positively influence prey rescue if predators preferentially consume maladapted individuals–accelerating the rate of prey adaptation–and the selective benefit outweighs the demographic cost (Jones 2008; Osmond et al. 2017). A similar concept also applies if maladapted individual experience greater interspecific competition (Osmond & de Mazancourt 2013). If selection and density-dependence act on prey birth rates, prey evolution may also be accelerated via the effect of predation to reduce prey density-dependence, reducing prey generation time (Osmond 2017; also see Draghi et al. 2024). Predators may also experience “indirect evolutionary rescue,” whereby rescue in a declining predator population is instead driven by prey adaptive evolution, if the prey exhibit trade-offs between growth and defense against predation (Yamamichi & Miner 2015; Cortez & Yamamichi 2019; Hermann & Becks 2022; van Velzen 2023). It is also possible for a species with lower population size to exhibit a higher probability of rescue than its more abundant competitor, if the resource distribution evolves towards its optimum and away from its competitor (Van Den Elzen et al. 2017).

Theoretical work has also suggested that negative density-dependence, particularly intra-specific competition, also reduces rescue probabilities (Chevin & Lande 2010, Osmond & de Mazancourt 2013, Nordstrom et al. 2023; Draghi et al. 2024), although the magnitude of this effect remains controversial in the literature (Bell 2017). While the models of negative density-dependence analyzed by these authors differ in certain mathematical specifications, all of them share a number of simplifying assumptions that obscure the relationship between demography and evolution. It is often either assumed that selection acts solely on intrinsic growth rates, *r*, (e.g. Chevin & Lande 2010; Nordstrom et al. 2023) or population carrying capacity, *K* (Osmond & de Mazancourt 2013), representing opposites on the continuum of *r*- and *K*-selection. However, these two parameters are likely not independent in reality (Pianka 1970, Mallet 2012, Travis et al. 2023), and there is much empirical evidence to suggest that *r* and *K* may co-vary positively or negatively due to varying life-history strategies in yeast (Wei & Zhang 2019), bacteria (Reding-Roman et al. 2017; Marshall et al. 2023), and spider mites (Bisschop et al. 2021). For example, selection acting to increase individual resource acquisition rates may increase *r*, but decrease *K* by reducing total resource availability among conspecifics at high densities (Agrawal & Whitlock 2012), thus intensifying density-dependence. However, selection acting to increase the efficiency of resource use would lead to a greater *r* as well as *K* by lowering the per-capita resource requirement for maintenance (Travis et al. 2023). In contrast, *r*-selected models of rescue assume that the intensity of negative density-dependence is solely a function of population size/density, and itself doesn’t evolve (since *K* is held constant), thus neglecting the above relationships. Currently, we lack a theoretical framework to understand evolutionary rescue through the evolution of ecological traits that differentially influence both *r* and *K*, representing a significant gap in the literature.

In this paper, we use a consumer-resource model with overlapping generations to investigate how evolutionary rescue of a consumer species is influenced by the ecological effects of an abrupt environmental change that, in the absence of evolution, would lead to extinction. Briefly, consumer-resource models explicitly track both consumer and resource population dynamics (see Sakarchi & Germain 2024 for a review). Consumer growth is a function of various ecological factors: resource acquisition rates, the efficiency of resource conversion into offspring, resource abundance, and mortality rates, whereas resource growth is typically modelled using logistic growth minus consumption by consumers. Consumer density-dependence arises via decreasing resource availability, settling on an equilibrium between resource consumption and replenishment. Using this framework, we characterize how evolutionary rescue in a consumer species is influenced by the ecological trait affected by the environmental change. We consider both a one-locus and polygenic basis to the affected ecological traits, exploring the interplay between demography, selection, resource competition, and generation time. We begin our analysis by deriving analytical predictions for the fixation probabilities of rescue alleles that influence each ecological trait, the expected number of rescue mutations to occur during the process of population decline, and the mean time to consumer extinction in the absence of evolution, which work well under weak selection and low individual turnover rates (i.e. slow-growing populations with long generation times). Given the complexity of our eco-evolutionary model, we then use forward-time simulations to explore areas of the parameter space where our theoretical approximations are not met, that is, for populations that exhibit rapid growth at low densities and have short generation times.

### Ecological model

We modelled the ecological dynamics of the system using a discrete-time consumer-resource model (MacArthur 1970; Abrams 2019). Consumer ecological dynamics are described by the following recurrence relation:

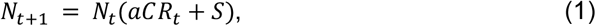

where *N*_*t*_ is consumer abundance at time-step *t, a* is the resource acquisition rate, *C* is the conversion efficiency of acquired resources into offspring, *R*_*t*_ is resource abundance at time-step *t*, and *S* is the survival rate. Our life-cycle proceeds with consumer births and resource consumption occurring simultaneously, followed by deaths. We consider overlapping generations using a simple form of age-structure. Individuals are able to reproduce and are susceptible to mortality beginning the time-step following their birth, and individual fecundity (*aCR*_*t*_) remains constant with age. An individual’s absolute fitness at time-step *t, W*_*t*_, is therefore *W*_*t*_ = *aCR*_*t*_ + *S*. The quantity *W*_*t*_ also describes the geometric growth rate of a lineage with trait values *a, C*, and *S*, in a population with resource abundance *R*_*t*_. The ecological dynamics of the resource are described by the following recurrence relation,

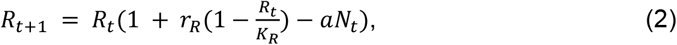

where *r*_*R*_ is the intrinsic growth rate of the resource and *K*_*R*_ is its carrying capacity.

### Simulation methods

We ran individual-based forward-time simulations using SLiM v4.01 (Haller & Messer 2022). Briefly, we initialized our simulations with consumers and resources at demographic equilibrium. An abrupt environmental change was introduced at time-step *t*_∗_ such that the population would go extinct in the absence of evolutionary rescue. A population was recorded as having gone extinct if its population size equaled 0. Conversely, a population was recorded as having rescued if it returned to a non-zero equilibrium, either through the fixation of a single beneficial allele (one-locus simulations) or through a return to its phenotypic optimum (quantitative trait simulations). Downstream analyses were performed using the programming language R 4.4.2 (R Core Team 2024).

We modelled the total number of births to be Poisson distributed with a mean of *N*_*t*_*aCR*_*t*_ for the consumers and a mean of 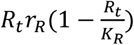 for the resources. Once the total number of offspring was established, parentage was retroactively assigned for consumer offspring by drawing from a multinomial distribution, with the probability of an individual being assigned parentage proportional to its fecundity. Consumer deaths were modelled by drawing from a binomial distribution with the probability of survival equal to the individual’s survivorship. We did not explicitly model resource death as its net growth is already captured by the *r*_*R*_ term, which we allocate to births for brevity. Resource consumption was modelled using a binomial distribution with *R*_*t*_ trials and probability of success *aN*_*t*_, the per-capita resource consumption rate.

### One locus rescue

Consider a monomorphic consumer population at demographic equilibrium, 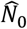 with ecological trait values *a*_0_, *C*_0_, and *S*_0_, consuming a resource population with intrinsic growth rate *r*_*R*_ and carrying capacity *K*_*R*_. An abrupt environmental change occurs at time *t*_∗_ reducing the mean growth rate of the resident population below unity (i.e.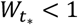) either via a reduction in individual resource acquisition rates, resource conversion efficiency, or survivorship. In the absence of evolution, the resident population, now exhibiting trait values *a*_res_, *C*_res_, and *S*_res_ will go extinct even at low density when the resource is at its carrying capacity (i.e., *a*_res_*C*_res_*K*_*R*_ + *S*_res_ < 1, such that 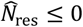, where 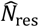 is the equilibrium consumer population size following the environmental change). Evolutionary rescue may occur if and only if a mutant lineage that may persist in this novel environment 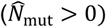 invades. Note that the mutant lineage need not return to 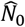. However, without loss of generality, we will assume in our simulations that all beneficial mutations have the effect of return the ecological trait to its value prior to the environmental change. We also assume for now that the consumer population is haploid and asexual to simplify the mathematics; we later relax this assumption when considering polygenic selection.

The probability of evolutionary rescue at time-step *t, P*_*t*_, is equal to one minus the probability that all rescue mutants arising at time-step *t* go extinct: 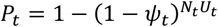, where *ψ*_*t*_ and *U*_*t*_ refer to the probability of fixation of a rescue mutation and the expected number of rescue mutations per individual at time-step *t*, respectively (Orr & Unckless 2008). Following this logic, the total probability of rescue is equal to 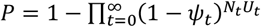. Assuming *ψ*_*t*_ and *N*_*t*_*U*_*t*_ are small for any given *t*, we approximate the above formula as an exponential decay:

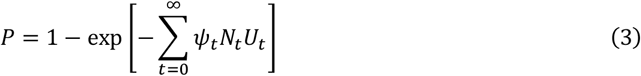

We solve *U*_*t*_ by considering that the number of mutations that arise at any time-point is simply a function of an individual’s fecundity multiplied by its mutation rate *μ*. Thus, the total mutation rate per individual at time-point *t* is

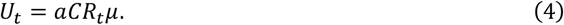

From eq. 4 we observe that the per-individual mutational supply will be decreased if an environmental perturbation reduces consumer fecundity instead of survivorship, all else remaining equal.

We now derive analytical predictions of *ψ*_*t*_ for mutant lineages. Consider a mutant allele that exhibits ecological trait values *a*_res_(1 + *δ*_*a*_), *C*_res_(1 + *δ*_*C*_), and *S*_res_(1 + *δ*_*S*_). The fixation probability of such a mutant allele will depend on its absolute fitness. If the individual first carrying the mutant allele has an absolute fitness below replacement (*a*_res_(1 + *δ*_*a*_)*C*_res_(1 + *δ*_*C*_)*R*_*t*_ + *S*_res_(1 + *δ*_*S*_) < 1), its lineage will deterministically go extinct. Fixation is only possible if the allele is able to increase its absolute representation in the population (*a*_res_(1 + *δ*_*a*_)*C*_res_(1 + *δ*_*C*_)*R*_*t*_ + *S*_res_(1 + *δ*_*S*_) > 1), although it must also escape stochastic loss when rare. Following the logic of branching processes, the probability that a new mutant allele experiences stochastic loss is equal to the probability that the individual in which it arises has some number of offspring, all of whose lineages eventually go extinct, and–with non-overlapping generations– that the individual itself fails to survive (Fisher 1922; Haldane 1927; Wright 1931; Otto & Whitlock 1997). The fixation probability of the mutant allele that occurs in a single individual at time *t* is thus equal to one minus its probability of stochastic loss. Assuming the number of offspring produced per individual in a time-step follows a Poisson distribution, we derive the following formula:

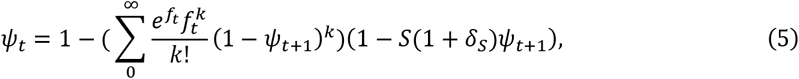

where *f*_*t*_ = *a*_res_(1 + *δ*_*a*_)*C*_res_(1 + *δ*_*C*_)*R*_*t*_ is the fecundity of a mutant individual at time-step *t*. Given that fecundity is density-dependent, *ψ*_*t*_ ≠ *ψ*_*t*+1_ in a population of changing size. Note that the fitness of the mutant allele is a function of the resource abundance, *R*_*t*_. Thus, in a declining consumer population (and increasing resource population), the individual carrying the mutant allele will experience an additional boost in fecundity and, consequently, an increased fixation probability the later its occurrence (i.e. *ψ*_*t*+1_ > *ψ*_*t*_). Assuming *a*_res_, *δ*_*a*_, *C*_res_, *δ*_*C*_, *S*_res_, *δ*_*S*_, and *ψ*_*t*_ are small and a separation of time-scales between consumer and resource growth, we derive a continuous-time approximation for the change in *ψ* as a function of *N*, which we solve to get the fixation probability as a function of the current population size, *ψ*(*N*_*t*_ ) (see *Appendix A* for the derivation). Ignoring terms of order O(*ψ*^2^) or above, the fixation probability of a mutant allele that increases fecundity for a given population size, *ψ*_*f*_(*N*_*t*_), either via increasing *a* or *C* is:

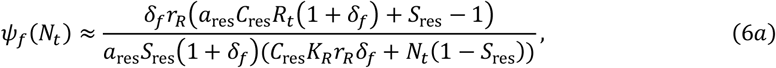

where *δ*_*f*_ = *δ*_*a*_ when the mutant allele increases *a* and *δ*_*f*_ = *δ*_*C*_ when it increases *C* (we do not consider the case where a mutant allele increases both). Conversely, a mutant allele that increases survivorship, *ψ*_*S*_(*N*_*t*_), will experience fixation probability:

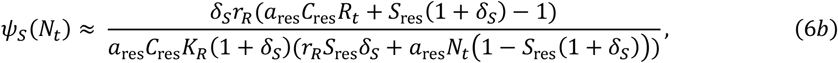

A quick analysis of eqs. 6a and 6b reveals that fixation probabilities of both fecundity and survival mutants are maximized as *N*_*t*_ → 0, as mutant individuals benefit from increased resource availability (*R*_*t*_ → *K*_*R*_), thus increasing their number of offspring.

Our last step is to derive a general solution for *N*_*t*_ and *R*_*t*_ in the resident population. To do so, we must make two key assumptions. First, we assume that resource abundance is at equilibrium with respect to consumer abundance and trait values (“separation of time-scales”) and, second, that few consumer birth and death events occurs per time-step such that consumer growth is approximately continuous (i.e. low individual turnover). Dropping subscripts, we find that 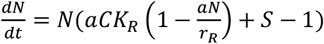, where 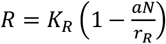. Here consumer population growth is logistic, and 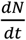 can be solved to get 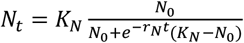, where *r*_*N*_ = *a C K*_*R*_ + *S* −1and 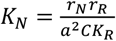. Note that 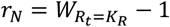, the growth rate of the consumer when resource availability is maximal. When *r*_*N*_ > 0, the population is able to grow at low density. This general solution for *N*_*t*_ can then be used to approximate *R*_*t*_ with the separation of time-scales assumption. We use these forms for *N*_*t*_ and *R*_*t*_ to estimate *ψ*_*t*_ and *U*_*t*_, from which we numerically estimate the total probability of rescue, *P*, using eq. 3. Mean times to extinction, *t*_*e*_, may be approximated by solving for the time-step at which there is only one consumer individual remaining in the population 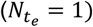 given the resident ecological trait values following the environmental change. Note that our quasi-deterministic approximation of *N*_*t*_ and *R*_*t*_ here will generally over-estimate *t*_*e*_ as it neglects demographic stochasticity and tends to over-estimate the resource abundance.

We also cross-examine our analytical predictions of *ψ*_*t*_ and *U*_*t*_ with numerical predictions. We numerically predict *ψ*_*t*_ using an iterative method whereby we solve for 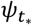 at equilibrium 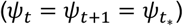, which occurs when *N*_*t*_ = *N*_*t*+1_ = 0. We then retroactively solve for *ψ*_*t*_ as a function of *ψ*_*t*+1_ (see eq. 4) by moving backwards in time from the point at which the consumer population went extinct. We numerically predict *U*_*t*_ by calculating it from observed consumer and resource abundances in the simulations, thus giving its expected value.

We use forward-time simulations to test the accuracy of our approximate theory and characterize how the probability of evolutionary rescue depends on whether acquisition rates, conversion efficiencies, or survivorships are negatively impacted. We simulate the scenario where an abrupt environmental change affects one ecological trait such that the population will go extinct in the absence of evolution, and we draw comparisons among these three scenarios with respect to their intrinsic growth rates following the environmental change, *r*_*N*∗_ (where *r*_*N*∗_ < 0). We model rescue mutations to occur *de novo* at constant rate *μ* per offspring following the environmental change, with rescue mutations returning the ecological trait to its value prior to environmentally induced maladaptation.

We simulate evolutionary rescue of a “slow-growing” consumer population (low fecundity, high survivorship) with an intrinsic growth rate of *r*_*N*_ = 0.03 and *K*_*N*_ = 2084 and a “fast-growing” consumer population (high fecundity, low survivorship) with an intrinsic growth rate of *r*_*N*_ = 0.4 and *K*_*N*_ = 2230 prior to the environmental change (see fig. 1 caption for parameters). Despite the number of assumptions made in our approximate theory, it performs remarkably well in the slow-growing case (Fig. 1A squares), when birth and death rates are low and selection weak. In the fast-growing case, the approximate theory substantially over-predicts rescue probabilities (Fig. 1B squares). Here our assumptions of small fixation probabilities and continuous-time population growth are not met due to strong selection and rapid consumer growth. This results in over-estimation of fixation probabilities and, more importantly, substantial over-estimates of the resource abundance and therefore mutational supply (see eq. 4). In this parameter space, consumer and resource intrinsic growth rates are the same order of magnitude, such that the resource does not reach a quasi-equilibrium with respect to consumer density.

**Figure 1.**
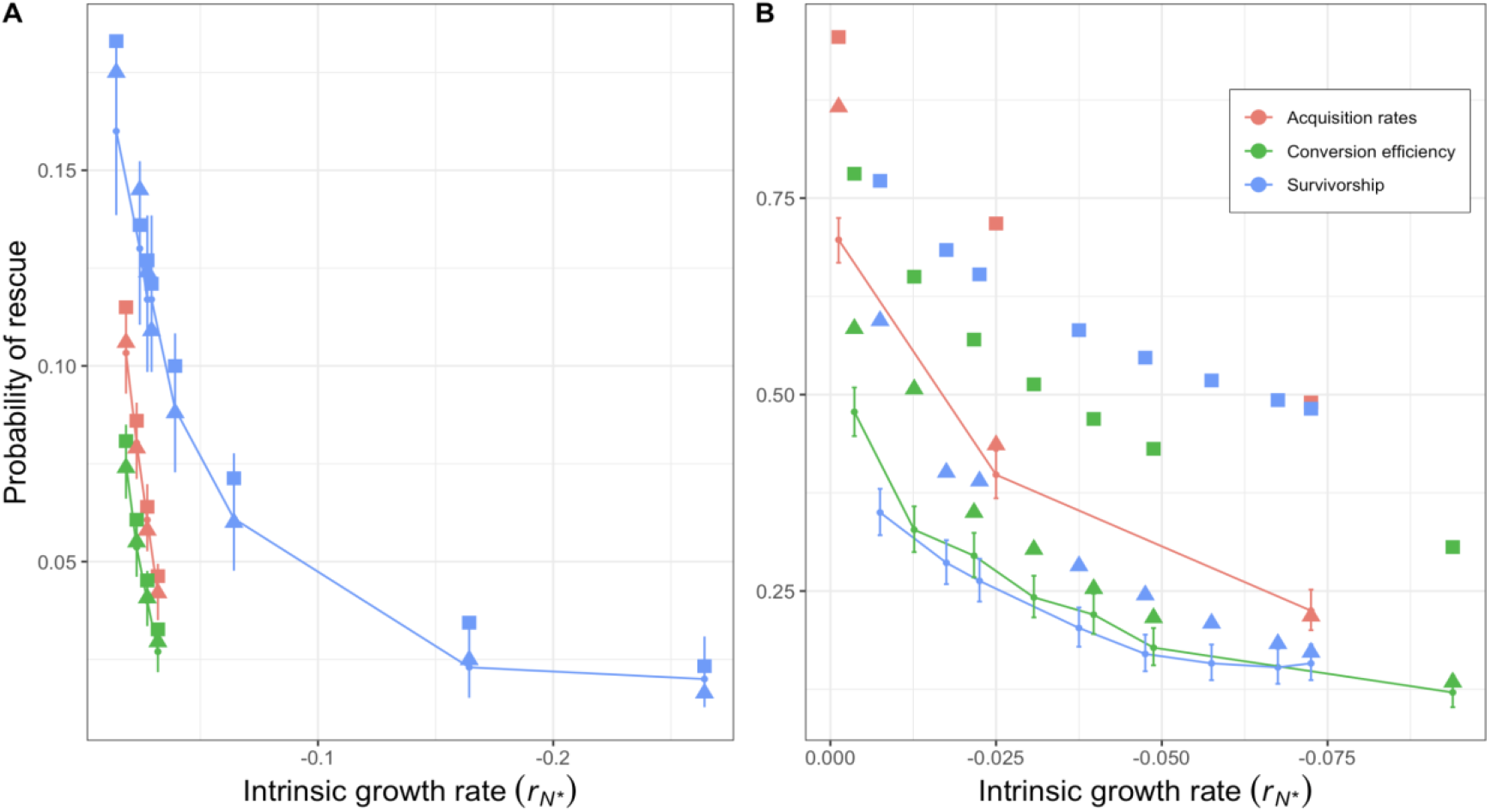
Probabilities of one-locus rescue as a function of population intrinsic growth rates and the affected ecological trait following an abrupt environmental change in (A) a slow-growing population (*a*_0_ = 9.5 × 10^−5^, *C*_0_ = 0.0095, *S*_0_ = 0.95, *r*_*R*_ = 0.475, *K*_*R*_ = 95,000) and (B) a fast-growing population (*a*_0_ = 9.5 × 10^−5^, *C*_0_ = 0.1 *S*_0_ = 0.5, *r*_*R*_ = 0.475, *K*_*R*_ = 95,000). The mutation rate is 10^−4^. Small dots connected by a line represent data from at least 1000 replicate simulations. Squares represent rescue predictions using eqs. 6a & 6b and our quasi-deterministic theory applied to eq. 3. Triangles represent numerical predictions of fixation probability using our iterative numerical method alongside estimates of expected mutation supply using simulation data.

We correct these discrepancies to some degree by instead using numerical estimates of fixation probabilities and estimated mutational supply from observed data, although we still observe some over-estimation of rescue probabilities for slower rates of decline (Fig. 1B triangles). In this regime, our theoretical framework will always over-predict fixation probabilities and ultimately rescue because it neglects several sources of stochasticity (birth, consumption, death) from our simulations that are most pertinent during intermediate and high consumer densities. Overall, our analytical theory provides a useful starting point to develop intuition for our rescue scenario, but is nonetheless limited in scope due to the complexity of our eco-evolutionary model, which incorporates aspects of density-dependent demography, selection, and overlapping generations with two demographic variables (*N*_*t*_ and *R*_*t*_).

The central result from our simulations is that the probability of evolutionary rescue depends on the affected (and subsequently evolving) ecological trait, and the ecological trait most favourable to rescue via *de novo* mutations depends on the population’s resident life-history (Fig. 1). Resident populations exhibiting the slow-growing strategy experience the greatest probability of rescue from declines in survivorship, followed by acquisition rates and then conversion efficiencies (Fig. 1A). On the other hand, populations exhibiting the fast-growing resident strategy exhibit the greatest probability of rescue following declines in acquisition rates, with declines in survivorship and conversion efficiency resulting in similar rescue probabilities (Fig. 1B).

When fecundity is negatively impacted, consumer populations always experience higher probabilities of rescue when their resource acquisition rates are reduced rather than their ability to convert acquired resources into offspring. Despite *a* and *C* mutants of equal effect size (*δ*_*a*_ = *δ*_*C*_ = *δ*_*f*_) experiencing equivalent fixation probabilities for any given consumer and resource density (eq. 6a), individuals carrying the rescue allele in a population experiencing reductions in acquisition rates will benefit from greater resource availability earlier on during the process of population decline (Fig. 2A), hastening its spread in the population (Fig. 2B).

**Figure 2.**
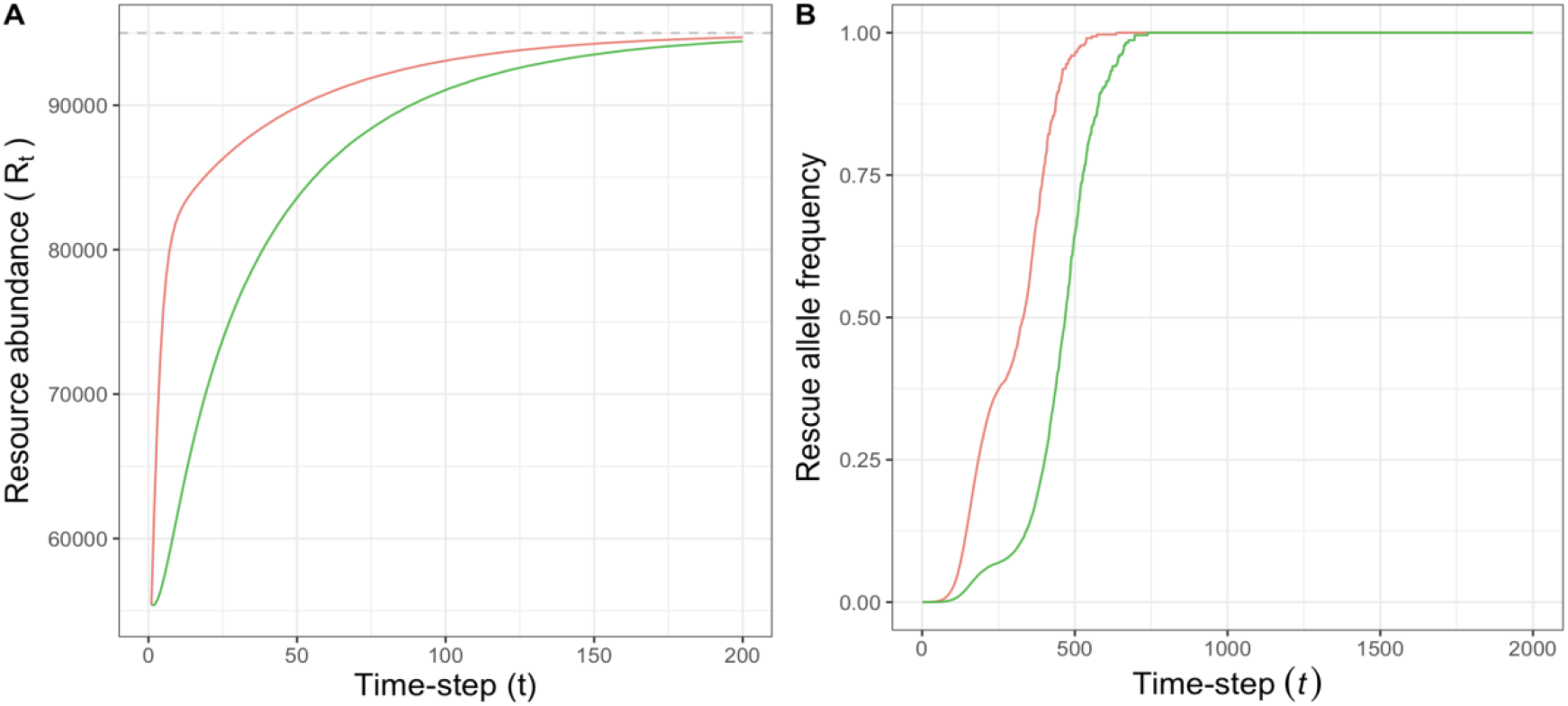
(A) Resource abundance and (B) rescue allele frequency as a function of time following an environmental change that reduces acquisition rates (orange) and conversion efficiencies (green). Observed data was averaged from 3000 replicate simulations of the slow-growing population with *a*_0_ = 9.5 × 10^−5^, *C*_0_ = 0.0095, *S*_0_ = 0.95, *r*_*R*_ = 0.475, *K*_*R*_ = 95,000, and *N*_0_ = 2084, where *a*_∗_ = 3.5 × 10^−5^ (orange) or *C*_∗_ = 0.0035 (green). In both cases, *r*_*N*∗_ = −0.018. The grey dashed line in (A) represents the resource’s carrying capacity, *K*_*R*_ = 95,000.

Whether rescue via *de novo* mutation is more favourable following declines in fecundity or survivorship is variable with respect to the population’s resident life-history strategy, and this result is primarily driven by differences in the supply of rescue mutations. Our simulation results suggest that rescue probabilities are approximately linear with respect to mutational supply (Figs. 3A & D) because we are only considering rescue via *de novo* mutations. In simulations of the slow-growing population, the affected ecological trait does not significantly influence time to extinction, *t*_*e*_ (Fig. 3C), although it is worth noting a slight discrepancy between the traits. Instead, decreases in survivorship result in a much greater total supply of rescue mutations than decreases in fecundity for any given *t*_*e*_ (Fig. 3B). Population mutation rates are a function of both total population size and birth rates. Thus, while all populations experience lower populational mutation input following the environmental change due to smaller population size, populations with reduced fecundity suffer additional reductions in mutation rates due to fewer births.

**Figure 3.**
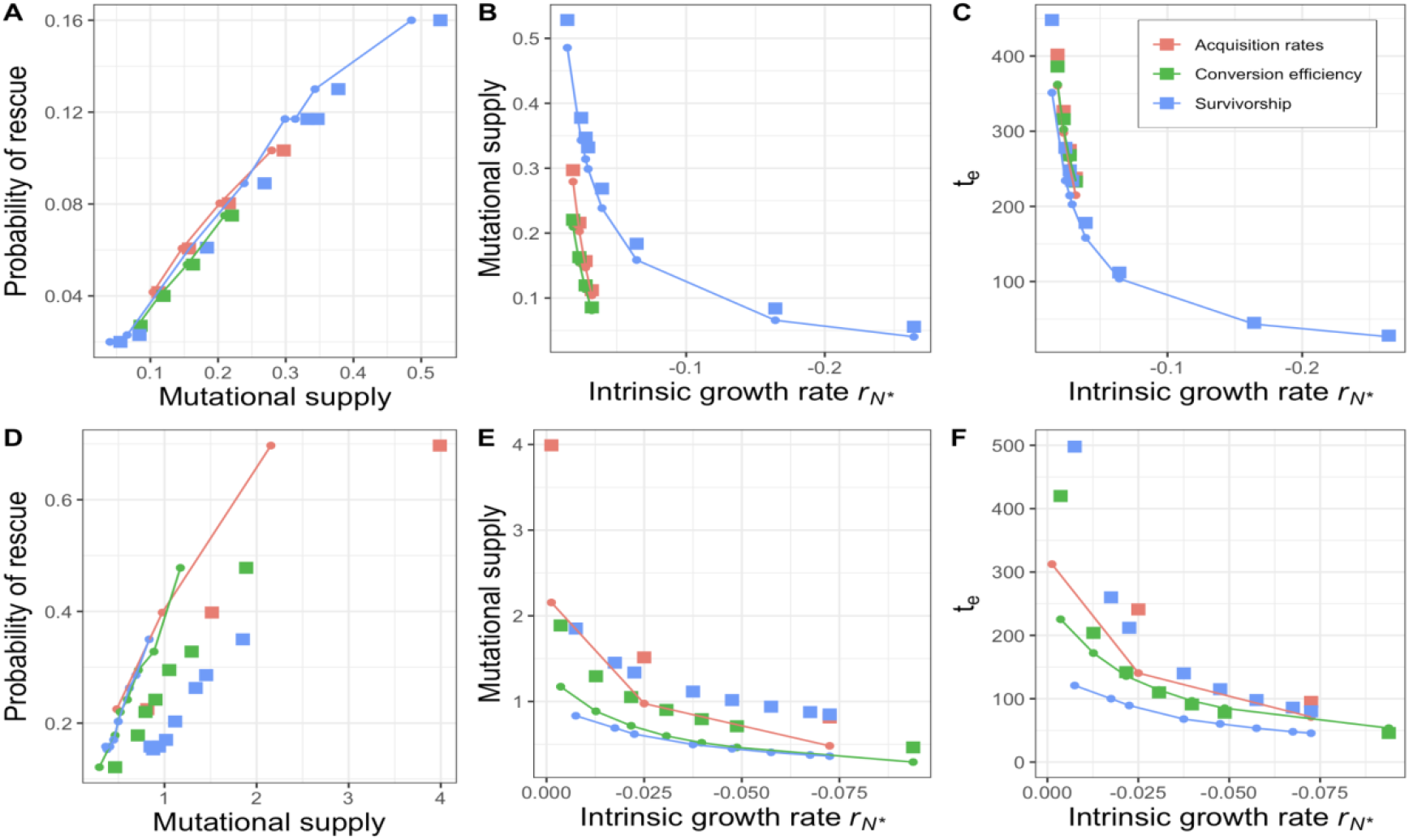
Probability of rescue, mutational supply, and times to extinction for a slow-growing population (A) – (C) and fast-growing population (D) – (F). The circles connected by lines represent observed mutation supply and times to extinction from at least 1000 replicate simulations, and the squares represent numerical estimates from our quasi-deterministic theory.

In the fast-growing population, on the other hand, it is following reductions in resource acquisition rates that the mutational supply, and therefore the probability of rescue, is maximized (Fig. 3D & E), and this result corresponds to a slower time to extinction (Fig. 2F). Interestingly, reductions in survivorship lead to a mutational supply almost indistinguishable, if not lower, from reductions in conversion efficiency (Fig. 3E). This result can be explained by considering the direct effect of survivorship on generation time, particularly since we model fecundity to be constant with age. The average generation time is 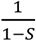, a function of survivorship. Reduced survival accelerates individual turnover and reduces generation time, thus leading to extinction in fewer time-steps than when fecundity is reduced, in the absence of evolution. This effect is marginal in the slow-growing population (where *S*_0,*slow*_ = 0.95), but occurs in much greater magnitude in the fast-growing population (*S*_0,*fast*_ = 0.5), where negative intrinsic growth rates via decreases in survivorship are only possible when *S*_∗,*fast*_ < 0.0975 (for *r*_*N*∗_ < 0), resulting in substantially reduced generation times and rapid individual turn over. The similar results between the survivorship and conversion efficiency cases are thus primarily driven by trade-offs in mutation supply per time-step and extinction time. These trade-offs also occur when comparing acquisition rates to survivorship, although populations with lessened acquisition rates additionally benefits from greater resource abundance at an earlier rate due to lower per-capita resource consumption rates, leading to the greatest probability of rescue here.

Although rescue via *de novo* mutation appears to be primarily driven by differences in total mutational supply, regardless of the ecological trait, it is worth briefly describing the mechanics of selection and fixation probabilities in our model, which indeed differ among ecological traits. An interesting result of our analysis is that the fixation probabilities, *ψ*, of rescue alleles at any time-point do not necessarily correspond to their selective advantage, *s*, defined as the fitness of individuals carrying the rescue allele relative to the declining resident allele (see Fig. 4 caption). Instead, the fixation probability of a rescue allele will be additionally dependent on the rate of consumer population decline as well as resource abundance, which are differentially influenced by the three ecological traits.

**Figure 4.**
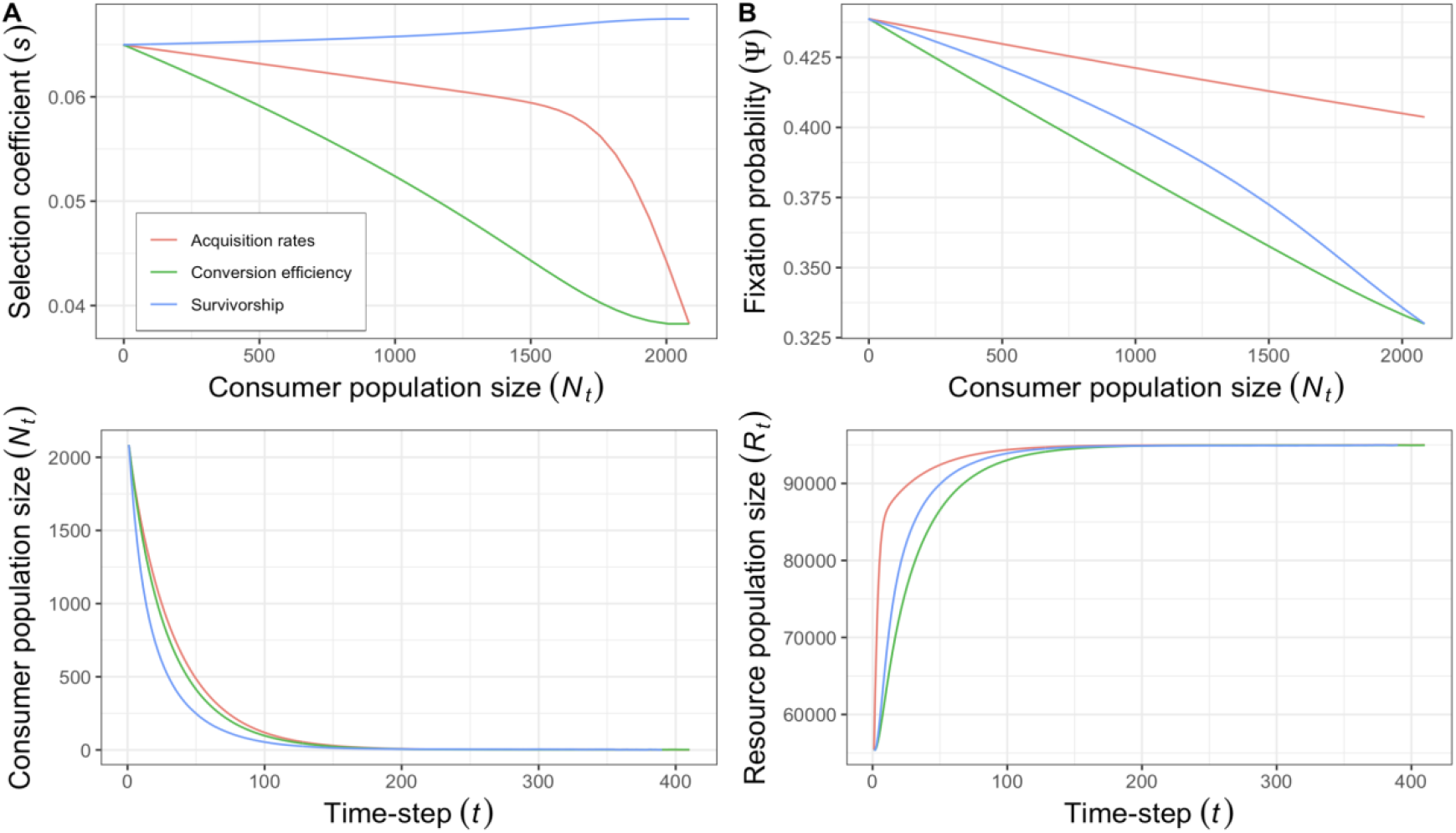
Selection coefficients, fixation probabilities, consumer and resource abundance over time in the slow-growing population. (A) Selection coefficients were calculated using the formula 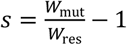, where *W*_mut_ is the fitness of the mutant type and *W*_res_ the fitness of the resident type in a population in need of rescue (*W*_res_ < 1). Data was averaged from at least 1000 replicate simulations of the slow-growing population with *r*_*N*∗_ = −0.027 from *a*_∗_ = 2.5 × 10^−5^ (orange), *C*_∗_ = 0.0025 (green), or *S*_∗_ = 0.8868 (blue). (B) Fixation probabilities were estimated using eqs. 6a and 6b, which perform well for these simulations (see Fig. 1A). (C–D) Mean consumer and resource abundance over time.

Selection on mutant alleles that increase fecundity, either via increasing *a* or *C*, are selectively favoured at low consumer density, whereas selection on increased survivorship is maximized at high consumer density (Fig. 4A). However, fixation probabilities of rescue mutants increase monotonically with declining consumer population sizes, albeit at different rates for each ecological trait, ultimately converging onto a single maximum as consumer population size declines towards extinction (Fig. 4B). This apparent contradiction between *s* and *ψ* may be resolved simply by considering that, despite simulations being matched for *r*_*N*∗_, such that their fixation probabilities once *N*_*t*_ = 0 are equal, they differ in their rates of decline and effects on resource abundance at intermediate densities (Fig. 4C & D). For example, when resource acquisition rates are reduced, the rate of consumer population decline is marginally slower, and the rate of resource growth marginally faster than a survivorship mutant, such that, despite having a lower selective advantage (*cf*. Fig. 4A blue & orange lines), a rescue allele that increases acquisition rates will experience a higher fixation probability at intermediate to high consumer population sizes. Conversely, despite decreases in survivorship leading to more rapid declines in consumer density than decreases in conversion efficiency (*cf*. Fig. 4A blue & green lines), the selective advantage of a survivorship mutant exceeds that of a conversion efficiency mutant, such that it experiences higher fixation probabilities for intermediate consumer population sizes (Fig. 4B).

### Polygenic rescue

Our analysis has until now been restricted to the case of evolutionary rescue via the *de novo* occurrence and fixation of a single beneficial allele in a haploid population, from which we develop some analytical theory and derive general intuition for the mechanisms underlying rescue. However, it is also possible, and indeed likely, that the genetic architecture underlying consumer ecological traits is polygenic, with standing genetic variation in ecological trait values prior to the environmental change. We explore this scenario by modelling a sexual, diploid consumer population with polygenic phenotypes underlying ecological performance. Our polygenic model slightly differs from those typically used in the classical quantitative genetics literature in that the phenotype instead determines an individual’s ecological trait value (*a, C*, or *S*), which then influences fitness (recall that *W* = *aCR*_*t*_ + *S*) instead of the phenotype directly influencing fitness (as in Gomulkiewicz & Holt 1995, Burger & Lynch 1995, etc.). Briefly, we consider an arbitrary phenotypic trait (e.g. height, size) of the consumer underlying a single ecological trait *ϕ* (where *ϕ* may refer to *a, C*, or *S*), holding all other traits constant. The phenotypic trait value *z* of an individual is determined by the sum of the genetic contribution of 100 loci with additive phenotypic effects plus a normally distributed environmental effect with mean 0 and variance 1. The ecological trait value of an individual with phenotypic trait value *z* is described by a Gaussian performance function:

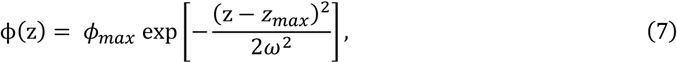

where *ϕ*_*max*_ refers to the maximum ecological trait value, *z*_*max*_ refers to the phenotypic value which maximizes ϕ (when z = *z*_*max*_, ϕ(z) = *ϕ*_*max*_ ), and *ω*^2^ describes the function’s width. Following a burn-in period where genetic variation is introduced via mutation-selection balance, we model rescue by imposing an abrupt environmental change that modifies *z*_*max*_ of one ecological trait to *z*_*max*∗_ such that *r*_*N*∗_ < 0 and the population will deterministically go extinct in the absence of evolution. The population must now evolve toward its new phenotypic optimum to increase ϕ, and therefore mean fitness, 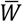, above one. At the time of the environmental change, the population experiences some additional reduction in fitness due to segregating variation around its optimum (variance load), but also experiences a head-start in the adaptive process following an environmental change due to some individuals carrying alleles that may move their phenotype towards the new optimum.

We again simulate the case of a slow-growing population 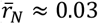, where the approximation is due to environmental variation in phenotype) with low fecundity relative to its survival, and a fast-growing population with high fecundity relative to its survival 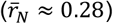, although with slightly different parameters. We recover a qualitatively similar result to our one-locus simulations, with rescue probabilities being contingent on both the affected ecological trait and the resident life-history of the population (Fig. 5). Decreases in survivorship are heavily favourable to rescue in slow-growing populations already exhibiting high survivorship, followed by acquisition rates and conversion efficiencies, a result almost identical to the one-locus case (*cf*. Fig. 1A & Fig. 5A). However, we observe some differences in the fast-growing case. Here, decreases in survivorship substantially hinder rescue, but rescue probabilities following decreases in acquisition rates or conversion efficiencies are similar.

**Figure 5.**
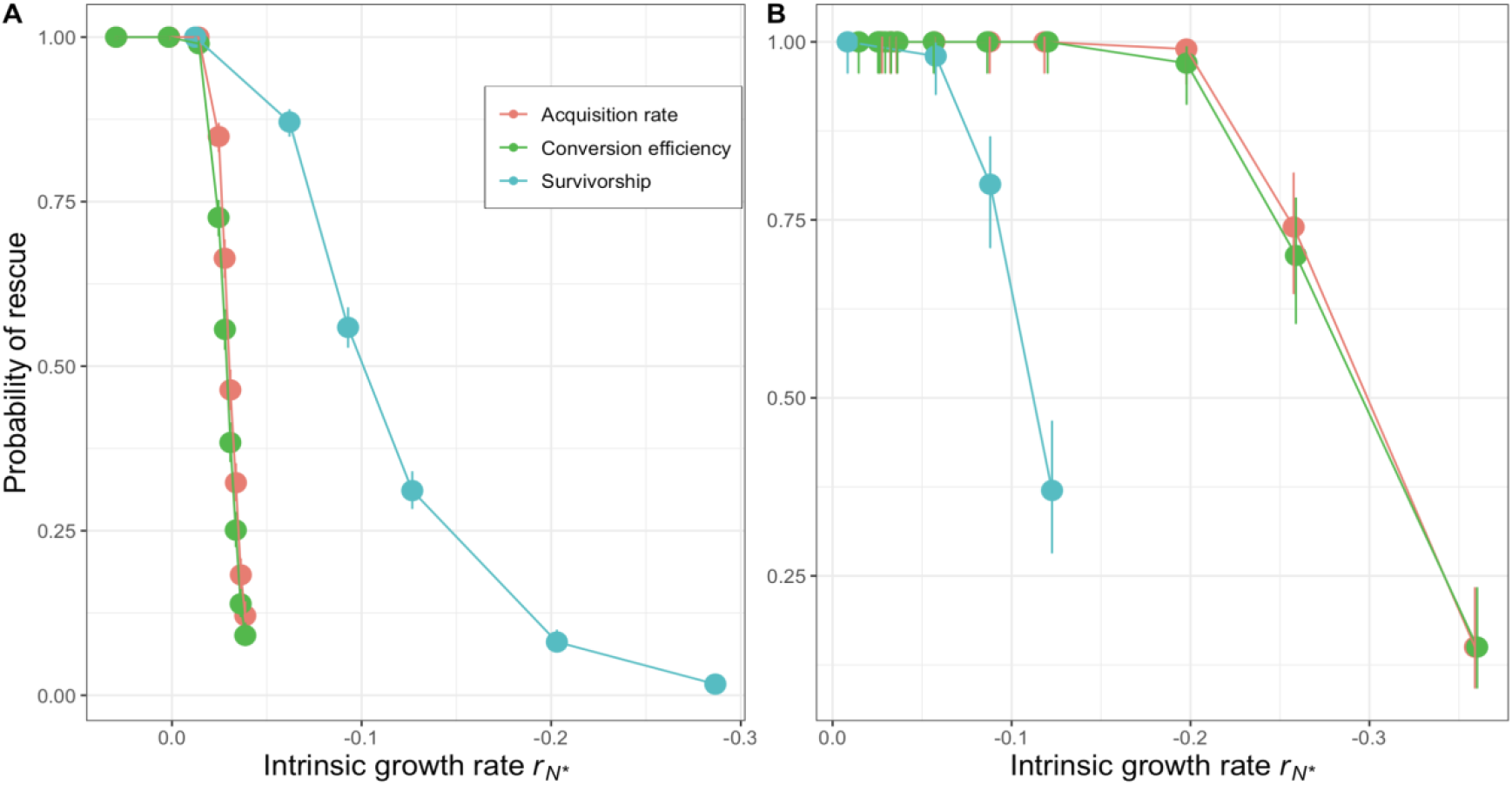
Rescue probabilities via evolution of a polygenic trait as a function of intrinsic growth rates and the affected ecological trait following an abrupt environmental change in (A) a slow-growing population (*a*_*max*_ = 10^−4^, *C*_*max*_ = 0.01, *S*_*max*_ = 0.99, *r*_*R,max*_ = 0.5, and *K*_*R,max*_ = 10^5^) and (B) fast-growing population (*a*_*max*_ = 10^−4^, *C*_*max*_ = 0.093, *S*_*max*_ = 0.5, *r*_*R,max*_ = 0.5, and *K*_*R,max*_ = 10^5^). The dots connected by lines represent observed data from at least 100 replicate simulations, and error bars represent 95% confidence intervals. The evolving ecological trait was determined by 100 loci evenly distributed among 10 chromosomes of map length 30cM. Mutations were drawn at per base-pair rate 10^−4^ from a normal distribution with mean 0 and variance 0.5.

In this scenario, *de novo* mutation plays a diminished role in the probability of rescue because of the occurrence of segregating variation. Thus, it follows that the advantage of greater mutational supply following decreased survivorship relative to decreased conversion efficiency is diminished, but populations with reduced conversion efficiency still benefit from slower times to extinction than populations with reduced survivorship (*cf*. Fig. 1B & 5B). Additionally, phenotypic change is no longer contingent on the establishment of large-effect beneficial mutations as in the one-locus case, but allele frequency change at many small effect loci resulting in the evolution of phenotypic trait values toward the new optimum. Thus, under a polygenic architecture, selective advantages experienced by favoured alleles play a relatively greater role in rescue than in the one-locus case due to the diminished role of mutational input.

### Selection

We now describe selection coefficients for mutant individuals relative to residents immediately following the environmental change and at low density. Immediately after an abrupt environmental change that would result in extinction of the resident type, the selection coefficient, *s*, experienced by a mutant individual is

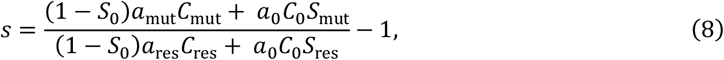

where *a*_0_, *C*_0_, and *S*_0_ are ecological trait values of the population prior to the environmental change, *a*_res_, *C*_res_, and *S*_res_ are ecological trait values of the resident allele following the environmental change, and *a*_mut_, *C*_mut_, and *S*_mut_ are ecological trait values of individuals carrying a mutant allele (see *Appendix B* for a derivation). From eq. 8 it is visible through the numerator that, at high density, the selective advantage of a fecundity mutant is proportional to 1 − *S*_0_, the mortality rate of the resident population prior to the environmental change, whereas the selective advantage of a survival mutant is proportional to *a*_0_*C*_0_, the total number of offspring produced per resource individual.

Consider the case where survivorship is reduced by an environmental change while fecundity remains unaffected, and there is the occurrence of a mutant allele that increases survivorship by factor *δ*_*S*_. In this case, *a*_mut_*C*_mut_ = *a*_res_*C*_res_ = *a*_0_*C*_0_, *S*_mut_ = *S*_res_(1 + *δ*_*S*_), and eq. 8 reduces to 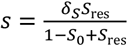. Note that this formula does not depend on fecundity-related traits and increases monotonically with *S*_0_. If instead an environmental change reduces fecundity, the mutant allele increases either acquisition rates or conversion efficiencies by factor *δ*_*f*_, and survivorship remains constant, eq. 8 reduces to 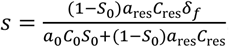, a monotonically decreasing function of *S*_0_. Thus, at high density, the selective advantage of a survivorship mutant will increase as a function of *S*_0_, whereas the selective advantage of a fecundity mutant will decrease as a function of *S*_0_. Additionally, given that the strength of selection is dependent on the evolving trait and resident life-history trait values, the equilibrium genetic variance available to the consumer population, due to mutation-selection balance, immediately following an abrupt environmental stressor will therefore also depend on the ecological trait under selection.

At low density, the selection coefficient is:

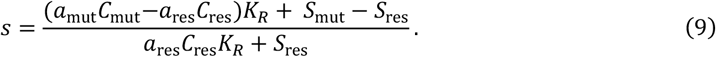

If selection acts on fecundity, then *a*_mut_*C*_mut_ = *a*_res_*C*_res_(1 + *δ*_*f*_) and *S*_mut_ = *S*_res_, eq. B4 reduces to 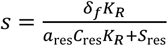. If we instead consider selection on survivorship *a*_mut_*C*_mut_= *a*_res_*C*_res_ and *S*_mut_ = *S* (1 + *δ* ), eq. B4 reduces to 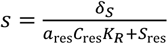. Thus, the selective benefit of a fecundity mutant will be scaled by the total resource availability at low density, such that its selective benefit will be orders of magnitude greater than that of a survivorship mutant at low density if their phenotypic effects are similar to one another (*δ*_*f*_ ≈ *δ*_*S*_). Additionally, at low density, selection does not depend on the ecological trait values of the population prior to the environmental change.

## Discussion

In this article we have shown that the probability of evolutionary rescue of a consumer species following an abrupt environmental change that reduces its intrinsic growth rate below zero is dependent on both the affected ecological trait and the population’s resident life-history strategy. In a slow-growing population, characterized by low fecundity and high survivorship, our analysis suggests that rescue is most favourable following reductions in survivorship, followed by resource acquisition rates and then conversion efficiencies. In a fast-growing population, characterized by high fecundity and low survivorship, it is instead following reductions in resource acquisition rates that rescue is most probable, followed by survivorship and resource conversion efficiency. These results are driven by the interplay between various factors, including the effects ecological traits have on the mutational supply, resource competition (i.e. density-dependence), selection, and times to extinction. We derive analytical formulae to predict the fixation probabilities of mutant alleles, rescue probabilities at any given time-point, and times to extinction, which work well when selection is weak and individual turnover rates are low.

There have been a number of previous studies on whether effects on either birth or death rates influence evolutionary rescue (Osmond et al. 2017, Czuppon et al. 2023, Raatz & Traulsen 2023, Draghi et al. 2024). In theoretical explorations of the evolution of antibiotic resistance, it has been suggested that rescue is most likely following increases in death rates versus birth rates due to reductions in competition experience by resistant strains (Czuppon et al. 2023; Raatz & Traulsen 2023) as well as increased mutational supply. Draghi et al. (2024) recover a similar result, finding that increased death rates lead to faster rates of adaptation, and therefore rescue, due to reduced generation time. In this paper we show that this is not necessarily the case; instead, rescue may be favoured following decreases in birth rates in populations with high fecundity. This is because, while increases in death rates may result in lower reductions in mutational supply per time-step and faster adaptation than reductions in birth rates, they also accelerate times to extinction, perhaps leading to a lower overall mutational supply. Moreover, the probability of rescue is affected by whether birth rates are reduced via reductions in resource acquisition rates versus their efficiency of conversion to offspring. In populations with already high fecundity and low survivorship, reductions in resource acquisition rates–but not resource conversion efficiency–lead to greater resource availability to rescue mutants, facilitating their establishment and therefore rescue to a greater degree than when mortality rates are increased.

Another novel aspect of our model is the analysis of the interplay between genetic architecture and the affected ecological trait on the probability of rescue in a two-species model. When rescue occurs via *de novo* mutation at a single locus, ecological variables such as resource availability and consumer vital rates substantially influence rescue through their effects on the establishment probability of beneficial mutations and the mutational supply. Conversely, when rescue occurs via both *de novo* mutation and standing genetic variation at many loci, differences in rescue more closely reflect differences in selective pressures, leading to a reduced benefit to increased mutation rates per time-step as in the case of increased mortality. Given that selection is indeed dependent on resident ecological trait values, this also suggests that the genetic variation available to selection for differing traits may indeed be dependent on the population’s resident life-history strategy, which may significantly influence rescue. For instance, Yamamichi et al. (2019) have shown, in a model of non-overlapping generations with a fluctuating environment, that increased survival may produce reservoirs of genetic variation (e.g. harbored by older or dormant individuals), although much of this variation would be deleterious due to those individuals phenotypically lagging behind fitter types. The analysis of evolutionary rescue under fluctuating environments instead of an abrupt environmental change may thus present an interesting extension to our work.

A central result of our simulations is that intrinsic growth rates alone are insufficient to predict the probability of rescue, because negative intrinsic growth rates may be the result of different ecological factors, each with a potentially different effect on factors such as the mutational supply, time to extinction, and density-dependence. To a first approximation, intrinsic growth rates may be a useful metric of population vulnerability, but our simulations suggest that the probability of rescue for a given intrinsic growth rate following an abrupt environmental change may be heavily dependent on the underlying ecological mechanism. However, it may be possible to predict these differences if empiricists are able to measure fecundity and viability directly.

One open question raised by our model may be what biological mechanisms may lead to decreases in traits such as acquisition rates, conversion efficiencies, and survivorship rates, and whether these have been observed. Regarding survivorship, there are various well-characterized examples of evolutionary rescue following antibiotic treatments in bacteria, herbicide treatment in weeds, and fungicide treatment for fungal diseases of economically important crops (see Bell 2017). Regarding acquisition rates and conversion efficiencies, Wei & Zhang (2019) show that quantitative trait loci (QTLs) in the yeast *S. cerevisiae* tend to show concordant effects on *r* and *K* (*r*/*K* trade-ups) in lower quality environments, but negative effects (*r*/*K* trade-offs) in higher-quality environments. They propose an explanatory model characterized by trade-offs between ATP production rates and utilization efficiency in the context of cell maintenance and division. Under low quality environments, *r* is low, and the evolution of increased *r* increases resource use efficiency, thus increasing both *r* and *K*. Conversely, in a high quality environment where *r* is already high, conversion efficiency decreases with increasing *r*. Here, the population is instead dominated by competition for the acquisition of resources. It is also well known that antibiotics may be distinguished into two classes according to mode of selection: drugs that act to increase mortality rates (bacteriocidal) versus decrease birth rates (bacteriostatic) (Czuppon et al. 2023). For instance, doxycycline, a bacteriostatic agent, acts by inhibiting protein production, and it has been shown that the evolution of doxycycline resistance in *E. coli* results in both increased intrinsic growth rates and carrying capacities (Reding-Roman et al. 2017). It may thus be possible to characterize the evolution of doxycycline resistance, and other chemically similar agents, in terms of the evolution of increased resource conversion efficiency. In general, empirical studies of evolutionary rescue often appear to focus on net rates of change rather than characterizing the specific ecological effects of environmental stressors. We thus highly encourage empiricists to pay greater attention to teasing out the effects of environmental stressors on birth and death rates when feasible, as this will lead to richer insights into the factors determining persistence in declining populations.

A major simplifying assumption of our theoretical analysis has been that ecological traits are uncorrelated, and only one trait is influenced by an abrupt environmental change and may subsequently evolve. This simple scenario is not intended to be a reflection of biological reality, but is a useful starting to point to characterize the core dynamics of the system. Survival and reproduction are indeed often correlated (e.g. Franco & Silvertown 2004), and evidence suggests that these correlations are altered by environmental change (Iles et al. 2019). We also do not consider a resource distribution, where competition for resource may lead to adaptive divergence among competing types (Dieckmann & Doebeli 1999). This would present an interesting extension to our model, as it is possible that adaptive divergence may relax competition, thus aiding rescue, but may result in less resource availability for divergent types, and therefore increased susceptibility to extinction over longer time-scales. Strong selective pressures imposed by an abrupt environmental change are also likely to bias the evolution of life-history traits to divergence between conspecifics, potentially promoting diversification (Chaparro-Pedraza & Bank 2025).

We have here provided an improved theoretical framework to understand how evolutionary rescue is influenced by the complex interplay between evolution and demography. The use of mechanistic models that provide increased biological realism is an important step forward in determining our treatment of populations at risk of extinction (Urban et al. 2024), an increasingly relevant problem given the rate of climate change (Berg 2010). The work presented here extends our understanding of the conditions under which we expect populations to adapt to environmental conditions that would otherwise lead to extinction.

## Appendix A. Derivation of *ψ*(*N*_*t*_)

Here we outline the derivation of the approximate fixation probability of a rescue mutant as a function of the current population size, *ψ*(*N*_*t*_), following Otto & Whitlock (1997). We evaluate the sum in eq. 5 from the main text to get:

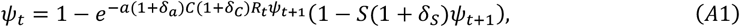

Assuming *a*(1 + *δ*_*a*_)*C*(1 + *δ*_*C*_)*R*_*t*_ and *ψ*_*t*+1_ are small, we perform a first-order Taylor series expansion of the exponential around 0 to get:

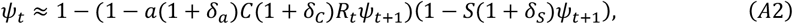

Since birth rates are density-dependent, and we are considering a scenario with variable demography, this is a time-inhomogeneous branching process (i.e. *ψ*_*t*_ ≠ *ψ*_*t*+1_). In a declining consumer population, for example, surviving individuals will benefit from greater resource availability in the future, leading to increased fecundity in future time-steps. Assuming the difference in fixation probabilities between any given time-step is small, we use a continuous-time approximation for Δ*ψ* = *ψ*_*t*+Δ*t*_ − *ψ*_*t*_. Substituting in the right hand side of eq. A2 in place of *ψ*_t_ and then taking the limit of 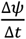 as Δ*t* → 0, we get 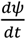. Dropping subscripts,

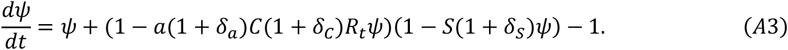

We can use the chain rule to re-write *ψ* as a function of population size 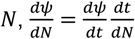. We derive 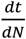 by assuming a separation of time-scales between consumer and resource growth. The logic behind this assumption is that if resource growth rates are much greater than consumer growth rates, then the resource will remain at a “quasi-equilibrium” state with respect to the consumer density at any time-step. Under these assumptions, consumer population dynamics are well represented by the logistic growth formula, 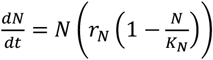 (Travis et al. 2023). We can solve for *r*_*N*_ by considering that the intrinsic growth rate of the consumer is simply its growth rate at low density, when the resource is at its carrying capacity. Specifically, when *N*_*t*_ = 0, *R*_*t*_ = *K*_*R*_ and *r*_*N*_ = *aCK*_*R*_ + *S* − 1. We find *K*_*N*_ by solving eq. 2 in the main text for 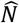, the consumer population’s demographic equilibrium. After some algebraic rearrangement, we get 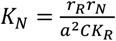. Plugging *r*_*N*_ and *K*_*N*_ into the logistic growth formula and simplifying, we get:

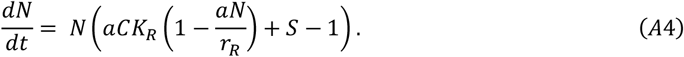

We can then solve for 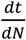 by taking the inverse of eq. A4:

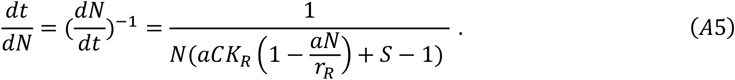

Note that 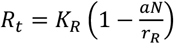, which can be found by solving eq. 2 in the main text for 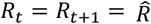. Since most individuals at any given time-point will be of the resident type, population dynamics will be a function of resident ecological trait values in a declining population. Combining eqs. A3 and A5 using the chain rule and re-writing *R* as a function of *N*,

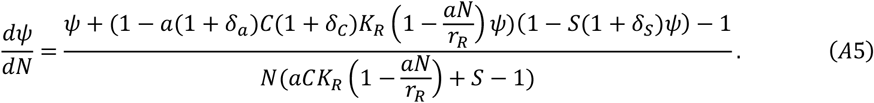

eq. A5 can be solved directly and simplified, depending on which ecological trait evolves, to get eqs. 6a and 6b.

## Appendix B. Selection at high and low density

Here we describe selection coefficients for a mutant allele that increases ecological trait values immediately following an abrupt environmental change that reduces fitness such that the consumer population will deterministically go extinct, and at low density, when the consumer population is nearly extinct. The fitness of the ancestral genotype after the environmental change is given by

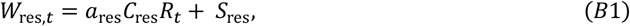

where *a*_res_, *C*_res_, and *S*_res_ refer to ecological trait values as changed by the new environment and *R*_*t*_ is the size of the resource population at time *t*. In order for the population to require evolutionary rescue, this fitness value must be less than one when the population is at low density; i.e., *a*_res_*C*_res_*K*_*R*_ + *S*_res_ < 1.

The invasion fitness of a rare mutant allele that occurs at time-step *t* is simply equal to its geometric growth rate,

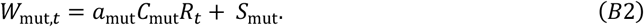

Therefore, the selection coefficient for the mutant allele at time *t* is given by

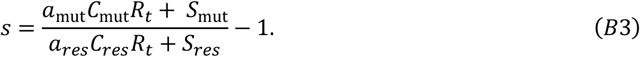

Immediately after the environmental change, resource abundance, *R*_0_, is still at equilibrium with respect to the ecological trait values prior to the environmental change, *a*_0_, *C*_0_, and *S*_0_, where 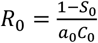 (which is derived from rearrangement of eq. 2 in the main text). Plugging this into our formula for *s* and after some rearrangement, we get

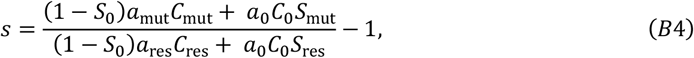

At low density, on the other hand, *R*_*t*_ = *K*_*R*_, from which eq. B2 simplifies to

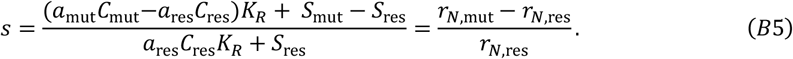

Here, the selective difference between the mutant and resident fecundity types is scaled by the carrying capacity of the resource.

## Notes

### Competing Interest Statement

The authors have declared no competing interest.

### Summary of Updates

The manuscript has been heavily revised to now include an explicit mathematical treatment of a one locus model of ecological trait evolution, now containing analytical approximations for the fixation probabilities of beneficial mutations, mutation supply, times to extinction, and the probability of rescue.

### Funding

- NSERC: RGPIN- 2016-03779, PGS D - 601214 - 2025

